# Pathogen priming alters host transmission potential and predictors of transmissibility in a wild songbird species

**DOI:** 10.1101/2024.10.21.619473

**Authors:** A.E. Leon, A. Fleming-Davies, J.S. Adelman, D.M. Hawley

## Abstract

Pathogen reinfections occur widely, but the extent to which reinfected hosts contribute to ongoing transmission is often unknown despite its implications for host-pathogen dynamics. House finches (*Haemorhous mexicanus*) acquire partial protection from initial exposure to the bacterial pathogen *Mycoplasma gallisepticum* (MG), with hosts readily reinfected with homologous or heterologous strains on short timescales. However, the extent to which reinfected hosts contribute to MG transmission has not been tested. We used three pathogen priming treatments– none, intermediate (repeated low-dose priming), or high (single high-dose priming)– to test how prior pathogen priming alters the likelihood of transmission to a cagemate during index bird reinfection with a homologous or heterologous MG strain. Relative to unprimed control hosts, the highest priming level strongly reduced maximum pathogen loads and transmission success of index birds during reinfections. Reinfections with the heterologous strain, previously shown to be more virulent and transmissible than the homologous strain used, resulted in higher pathogen loads within high-primed index birds, and showed higher overall transmission success regardless of host priming treatment. This suggests that inherent differences in strain transmissibility are maintained in primed hosts, leading to the potential for ongoing transmission during reinfections. Finally, among individuals, transmission was most likely from hosts harboring higher within-host pathogen loads, while associations between disease severity and transmission probability were dependent on a given bird’s priming treatment. Overall, our results indicate that reinfections can result in ongoing transmission, particularly where reinfections result from heterologous and highly transmissible strains, with key implications for virulence evolution.

**Importance:** As Covid-19 dramatically illustrated, humans and other animals can become infected with the same pathogen multiple times. Because individuals already have defenses against pathogen their immune systems have encountered before, reinfections are typically less severe, and are thought to be less contagious, but this is rarely directly tested. We used a songbird species and two strains of its common bacterial pathogen to study how contagious hosts are when their immune systems have some degree of prior experience with a pathogen. We found that reinfected hosts are not as contagious as initially infected ones. However, the more transmissible of the two strains, which also causes more harm to its hosts, was able to multiply more readily than the other strain within reinfected hosts, and was more contagious in both reinfected and first-infected hosts. This suggests that reinfections might favor more harmful pathogen strains that are better able to overcome immune defenses.

## Introduction

Reinfections are a common but understudied feature of many host-pathogen systems (1–4), including those of humans (e.g., SARS-Cov-2; (5)). The apparent pervasiveness of reinfections is somewhat surprising given that vertebrate immune systems harbor specific immune memory (6), allowing hosts to respond more rapidly and effectively to reinfection with the same pathogen. Nonetheless, the immune memory generated by prior pathogen infection is often incomplete, meaning some degree of reinfection is possible in many systems (7–9). The extent to which acquired protection from infection is “incomplete” can also increase over time as an individual’s initial acquired immunity wanes, increasing the risk of reinfection (10, 11). Overall, the growing recognition that many vertebrate host-pathogen systems are characterized by reinfection potential, whether immediately following recovery from initial infection or after initial acquired protection has waned, has led to recent calls for epidemiological models that better account for variability in infection-derived immunity (reviewed in 9). Nonetheless, the extent to which reinfected hosts contribute to ongoing pathogen transmission is still not well understood, despite the importance of this question for both epidemiological and evolutionary dynamics of host-pathogen interactions (8, 12).

Because initial pathogen infection generates acquired immunity with some degree of specificity for many hosts, reinfections with the same pathogen generally result in lower pathogen loads, reduced disease severity, and/or increased survival relative to hosts infected for the first time, which have no acquired protection (e.g. 13). Lower pathogen loads during reinfection are predicted to reduce a reinfected host’s transmission potential relative to a host infected for the first time. In two studies that experimentally infected mice with *Plasmodium chabaudi* parasites and then rapidly treated them to create “immunized” mice, a three to four-fold reduction in the density of transmission stage parasites was documented in previously immunized versus non-immunized mice (14, 15). Interestingly, the extent of reductions in transmission-stage parasites due to prior immunization was equivalent for homologous versus heterologous challenge strains of *P. chabaudi*, though heterologous strains were better able to transmit to mosquitoes (14). Further, numerous studies of vaccinated hosts across diverse taxa find that hosts challenged with the specific pathogen they were vaccinated against show lower transmission ability relative to unvaccinated hosts (16–19). Notably though, effects of vaccination on host transmission probability can also vary across pathogen strains. For example, vaccination of chickens for Marek’s virus reduced their transmission potential (relative to unvaccinated hosts) for a low-virulence strain of virus, but actually enhanced transmission potential of high-virulence strains (17). This occurred because vaccination against Marek’s virus generates incomplete immunity, protecting hosts from viral-induced mortality but not viral replication; together, this extends the infectious periods for virulent strains in vaccinated versus unvaccinated hosts, the latter of which rapidly succumb to virulent strains, often prior to transmitting (17). Thus, in addition to host pathogen loads, it is important to understand how immunization from prior infection or vaccination influences disease severity, which may determine reinfected host survival.

Other characteristics of pathogen strains, in addition to virulence, can potentially influence transmission potential during reinfection. Across taxa, inherent differences in strain within-host replication rates are often positively associated with transmission rates (20), suggesting that strain characteristics that influence within-host replication can, at least in some cases, predict transmission potential. High within-host replication rates are, in turn, associated with strain virulence in many systems (20, 21); for example, across ten parasite clones of *P. chabaudii* infecting laboratory mice, parasite clone growth, virulence, and transmissibility were positively related (15). Although strains infecting immunized mice showed overall reductions in all three pathogen fitness traits, the positive relationships between clonal growth, virulence, and transmissibility persisted in immunized hosts, potentially favoring virulent strains able to generate sufficient within-host growth, and thus transmissibility, in immunized hosts (22, 23). In addition to traits such as virulence and transmissibility, antigenic relationships among strains can determine the ability to sufficiently overcome host immune protection generated by initial infection: for example, serum from humans previously infected with a variant of SARS-CoV-2 showed stronger neutralization ability against homologous versus heterologous viral variants (24). Overall, such studies suggest that transmission success during reinfection can be strain specific, with transmission more likely during reinfections with heterologous and/or more virulent strains (14, 25, 26), provided such strains can better escape or overcome the acquired protection present in immunized hosts (22).

In addition to strain characteristics, the extent to which reinfected hosts transmit is likely dependent on the strength of acquired protection harbored by an individual at the time of reinfection. For example, the degree of SARS-Cov2 infectiousness (viral load) during reinfections or breakthrough infections was lowest for individuals who had both been vaccinated and experienced prior natural infection (27), and for malaria, individuals with prior infections in quick succession had serum stronger transmission-blocking immunity(28). Natural host-pathogen systems are inherently variable in the extent of initial pathogen exposure that hosts experience (e.g. 29, 30). Given that the strength of protection acquired from initial infection can vary with the dose (28, 29, 31) and frequency (28, 32) of prior pathogen exposure that a host experiences, variation in the extent of pathogen priming is predicted to influence the likelihood of ongoing transmission during reinfection.

Natural systems in which reinfections are common, such as the bacterial pathogen *Mycoplasma gallisepticum* (hereafter “MG”) of house finches (*Haemorhous mexicanus*), allow examination of how the extent of pathogen priming alters host transmission potential during reinfection with distinct pathogen strains (e.g., homologous versus heterologous). MG causes seasonal epidemics of mycoplasmal conjunctivitis in house finch populations (33). MG is largely transmitted at bird feeders (34), and because this obligate pathogen is short-lived outside of the host (35), finches experience variable levels of exposure at contaminated feeders. Diseased finches in the wild recover at high rates (36), and experimental studies show that reinfections are characterized by significantly lower pathogen loads and disease severity relative to first infections (37). Nonetheless, recovered individuals remain susceptible to reinfection with both homologous and heterologous strains (8, 37, 38), with little evidence for additional protection associated with reinfection strain homology (8).

While prior work suggests that reinfections are common in this host-pathogen system, the likelihood and severity of reinfection varies with both the degree of initial pathogen priming and the identity of the reinfecting strain. Leon and Hawley (39) experimentally varied the degree of pathogen priming experienced by finches, finding that a single high-dose MG priming treatment results in stronger host protection from homologous reinfection than intermediate degrees of MG priming such as repeated, low-dose exposures. A follow-up study (40) using similar priming treatments three distinct MG strains found that reinfections of primed hosts by a heterologous, more virulent MG strain were associated with higher within-host pathogen loads relative to MG strains with lower virulence and within-host replication rates (8, 40). Because within-host pathogen loads serve as a potential proxy for transmission likelihood, such results suggest that reinfection with a heterologous, more-virulent strain may result in higher transmission in this system, but work to date has not directly assessed transmission success. Given that prior studies have found discrepancies between effects of host immunization on proxies of transmission (such as the density of transmission-stage parasites) versus direct measures of transmission success (e.g. 14), it is key to examine effects of pathogen priming on between-host transmission success to fully uncover the importance of reinfections for pathogen ecology and evolution.

While no studies have examined MG transmission potential during host reinfection, several past studies quantified transmission in immunologically-naïve house finches. Among MG strains, higher within-host pathogen loads are associated with higher transmission rates (41), within a given strain, higher pathogen loads result in greater deposition of MG onto feeder port surfaces (42). In addition to pathogen loads, host disease severity is associated with transmission potential (43), with birds with more severely inflamed conjunctiva more likely to transmit to cagemates, even when accounting for the higher pathogen loads associated with higher disease severity (44–46). Given that reinfections in this system (8, 39) and others (15) are characterized by significant reductions in disease severity relative to hosts infected for the first time, the respective roles of pathogen load and disease severity in predicting transmission potential for pathogen-naive and reinfected hosts is key for understanding the selective pressures on pathogens to cause higher disease severity (i.e. virulence) in hosts.

Here we test how pathogen priming alters transmission potential for hosts reinfected with one of two strains (homologous versus heterologous), and quantify individual correlates of transmission success (disease severity, pathogen load). Although we did not have sufficient strain replication to isolate effects of strain virulence *per se* on transmission success, our reinfections used two strains that differ in virulence (8, 47), within-host replication rate (40, 47) and transmission potential (41). We specifically selected a more virulent strain as our heterologous strain for reinfections because MG strains collected from free-living house finches have increased in virulence over time (47, 48), with more virulent strains associated with higher transmissibility (41, 44). Thus, our experimental design mimicked a natural population in which individuals are most likely to be reinfected by either an endemic strain homologous to that which the host recently recovered from, or by an invading heterologous strain that has higher inherent transmissibility and virulence (44). To create variation in the degree of priming experienced by hosts, we varied the number and concentration of priming doses with a single MG strain (VA94) to create three priming levels: none, intermediate, or high. After recovery, we (re)-inoculated hosts and assessed effects of priming treatment on transmission success to a naïve cagemate during reinfection with one of two MG strains (the homologous strain, VA94, or the heterologous strain, NC06). Based on prior work (39, 40), we predicted the highest priming level would result in lowest host transmission potential. We also predicted that, consistent with prior work (40, 41), the heterologous, more virulent strain (NC06) would have higher overall transmission success, regardless of host priming treatment, with potential interactive effects of host priming and strain identity on transmission, as detected in the rodent-malaria system (e.g. 14). Finally, based on prior work (44, 45), we predicted that disease severity would correlate with transmission potential in naive hosts, but less so for reinfected hosts, which typically show stronger protection from disease versus pathogen loads (39).

## Methods

### Experimental Design

Seventy-eight captive, MG-naïve house finches (see *Supplement*) were randomly assigned to one of three priming treatments (n=26/group) that varied in both dose of pathogen exposure and total number of exposures (Figure 1). These priming levels - a negative control group (no priming), a low-dose repeat exposure group (“intermediate” priming), and a single high-dose exposure group (“high” priming) - were selected because they produced the greatest range in acquired protection in prior work, as measured by pathogen loads and disease severity during reinfection challenge (40). Inoculations for the intermediate priming group, which received six sequential exposures at 10^1^ color changing units (CCU)/ml, were given every other day and designed so that all groups received their final inoculation on the same day to allow for the same window of recovery prior to challenge (Fig. 1).

**Figure 1.**
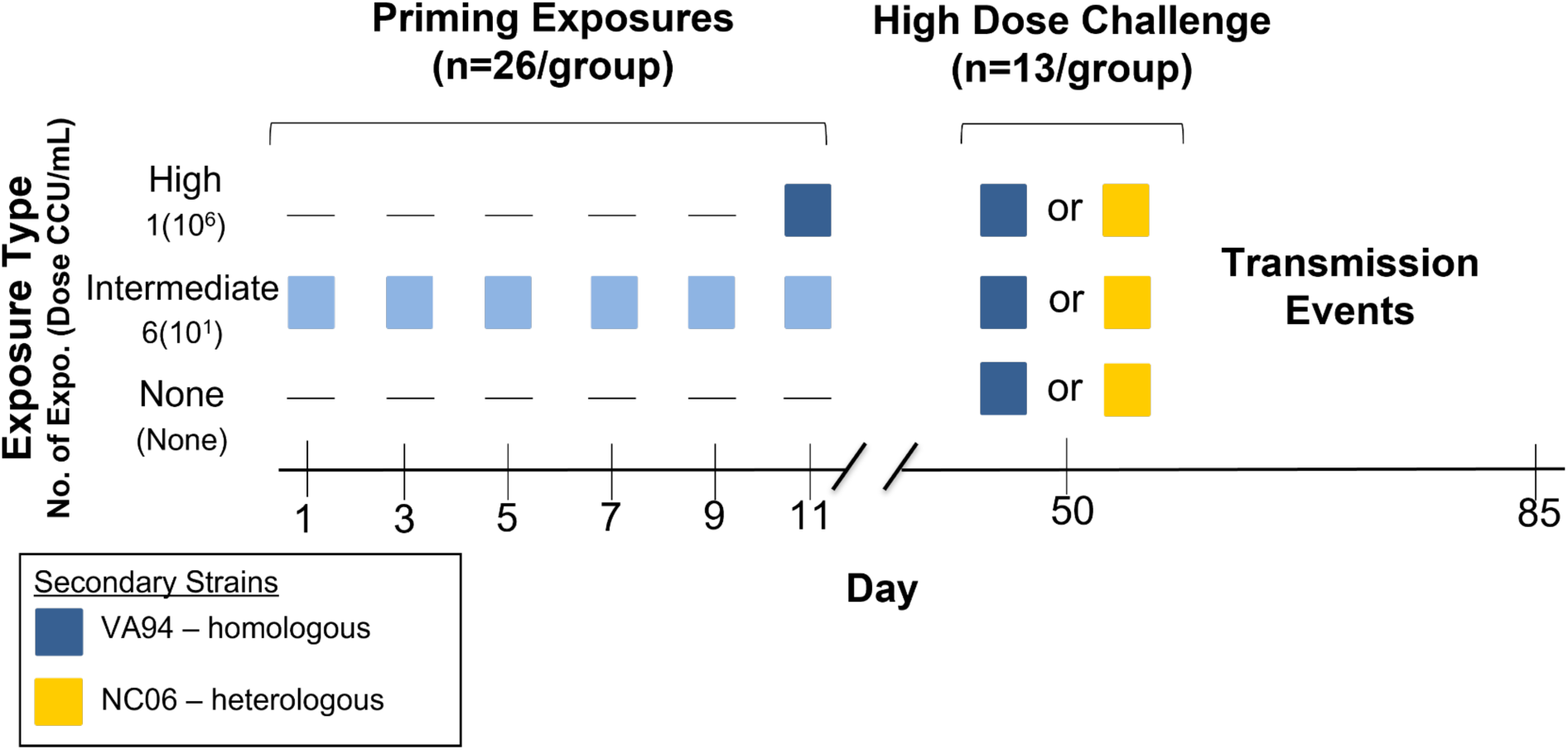
Experimental design and timeline. Index individuals (housed alone for the priming portion) were given one of three priming treatments (y-axis) with the *M. gallisepticum* strain VA94 (blue squares), with treatments varying by number of priming exposures and dose (the color gradient indicates variation in dose, with more intense color indicating increasing dose concentration; CCU = color changing units). On day 39 post-priming treatment, all index birds challenged with one of two strains: one homologous to that used in priming exposures (VA94, seen in blue) or a heterologous strain (NC06, seen in yellow) and then pair-housed with an MG-naive cagemate to assess pairwise transmission success.

Individuals were housed alone during priming treatment and given 39 days to recover from priming exposures, consistent with prior work (40). All birds then received a secondary, high-dose challenge with either a homologous or heterologous strain of MG (n=13 pairs/group; Fig. 1). At the time of secondary challenge, each of these 78 individuals was pair-housed with an MG-naïve cagemate to determine how pairwise transmission success varies with both the degree of pathogen priming and strain identity. Hereafter, individuals who acted as “transmitters” are referred to as “index” birds, and their immunologically-naïve cagemates used to assess transmission potential are referred to as “naïve” birds. Sex ratios of index birds were even for priming treatment groups (13:13 male:female). For secondary treatments, all transmission pairs were same sex, but because groups had a sample size of 13 (Fig 1), sex ratios were randomly assigned as either 6:7 male:female pairs or 7:6 female:male pairs. All protocols for animal use were approved by Virginia Tech’s Institutional Animal Care and Use Committee. See *Supplement* for capture and housing details.

### Pathogen inoculations

Two strains were selected based on previous work demonstrating differences in the maximum and average pathogen loads and virulence they produce in immunologically-naïve house finches (8, 47). The house finch MG strain “VA94” (7994-1 7P 2/12/09) (49) was used for all priming exposures, whether intermediate or high priming (Fig 1). Secondary challenge inoculations were all at high dose and varied only in strain identity: either the priming strain (homologous) or a heterologous strain “NC06” (2006.080-5 (4P) 7/26/12), used to represent a hypothetical invading strain with higher transmissibility. Inoculations were administered via droplet installation directly into the conjunctiva (70uL total volume across both conjunctiva) via micropipette (see *Supplement*). The negative control (sham inoculation) group received 70uL of sterile media.

### Disease severity and pathogen load

Disease severity was assessed by scoring the degree of visible inflammation, eversion, and exudate in conjunctival tissue (50) on a scale of 0-3 for each eye, and summing across eyes per individual within a given sampling date. Scoring was done blind to treatment. Pathogen load was assessed via swabbing of conjunctival tissue and MG-specific qPCR (see Supplement), with loads log10 transformed for analysis.

Index birds were eye scored and sampled for pathogen load on post-secondary inoculation days (PSID) 4, 7, 14, 21 and 28. To obtain high resolution data on transmission timing, naïve cagemates were eye scored daily on PSID 5 through 18 and then on days 21, 23, 25, 28, 32 and 35. Additionally, to ensure all individuals were still naive to MG just prior to the start of the experiment, we sampled for eye score and pathogen load on pre-inoculation day 19, as well as pre-challenge day 4 to obtain baseline data prior to re-inoculation. Responses to priming exposure levels have been previously examined (39), and are not included here.

### Transmission

Pairwise transmission was quantified as successful when a previously naïve cagemate developed scorable eye lesions (>0). Although the use of eye score as the assay for transmission can miss low-level, subclinical infections, prior work comparing the two metrics (34) showed that using eye score as the transmission metric robustly captures naïve individuals with minimum pathogen loads (> 1349 copies across both conjunctiva) considered to be infectious in this system. Further, the use of eye score eliminates potential false positives, which are known to occur in our qPCR assay (39).

### Analyses

All analyses were done using the statistical software R (51). In models where interactions were not significant, overall level effects were analyzed using Type II Likelihood Ratio tests using the car package in R (52). Whenever significant interactions were present, effects were analyzed using a Type III Likelihood Ratio (52). Post-hoc pairwise differences for significant interactions of interest were generated using the emmeans function and a Tukey adjustment for multiple comparisons.

#### Within-Host Responses

Because we were interested in the extent to which transmission success was a function of within-host responses, we analyzed both disease severity and pathogen load during secondary challenge of index birds with distinct priming treatments. As in our prior work (8), we ensured independence of repeated-measures data by analyzing only the maximum eye score and pathogen load for each index bird across four post-secondary challenge time-points (PSID 7, 14, 21, 28). For both models, fixed effects included priming treatment, secondary strain, and an interaction between the two effects (removed if not significant). To account for the non-continuous nature of score data, maximum eye score was treated as an ordinal factor and analyzed using cumulative link models (CLM) in the ordinal package (53). Maximum pathogen load was analyzed using a generalized linear model with a Gamma distribution, chosen because pathogen load was positively skewed, and inverse link function (lme4 package in R, (54)).

#### Transmission

Pairwise transmission (Y or N), assessed via any visible eye lesions in cagemates, was analyzed using logistic regression with binomial distribution and a logit link function. Fixed effects included priming treatment and secondary challenge strain. An interaction between priming exposure and secondary strain was tested but not included in the final model, as we did not have sufficient statistical power to fit a model with interactions.

To determine which host factors (eye score, pathogen load, or both) are predictive of transmission success across priming treatments, pairwise transmission was also analyzed across individuals using a second logistic regression with binomial distribution and a logit link function. Because our analyses of eye score and pathogen load indicated that priming treatment influences each host response somewhat distinctly, we included priming treatment in interaction with maximum pathogen load and eye score in the model. Although previous work found correlations between pathogen load and eye score across MG strains (47), in this study, the two variables were correlated at a level of 0.48 (Pearson correlation coefficient) across individuals, allowing us to include these variables as independent fixed effects in our model (55).

## Results

### Index Bird Within-Host Responses

All birds recovered from priming-induced disease (eye scores = 0) by the time of reinfection challenge on day 39. The level of pathogen priming that an index bird received significantly predicted host disease severity (i.e., maximum eye scores) during reinfection challenge, with the lowest disease in birds that received high pathogen priming prior to challenge (CLM estimates: intermediate priming: −1.48 ± 0.71; high-dose priming: −3.34 ± 0.93); priming treatment LR Chisq= 31.8, df = 2, P <0.0001; Fig. 2A). As expected, the more virulent NC06 strain produced higher maximum disease severity in birds than VA94 (strain[NC06]:1.59 ± 0.72), but secondary strain was not significant in the overall model (secondary strain LR = 2.15, df = 1, P = 0.14). There was no significant interaction between priming treatment and secondary strain identity on disease severity in index birds (priming:secondary LR = 5.10, df = 2, P = 0.078), suggesting effects of priming on disease severity were largely similar between the strains.

**Figure 2.**
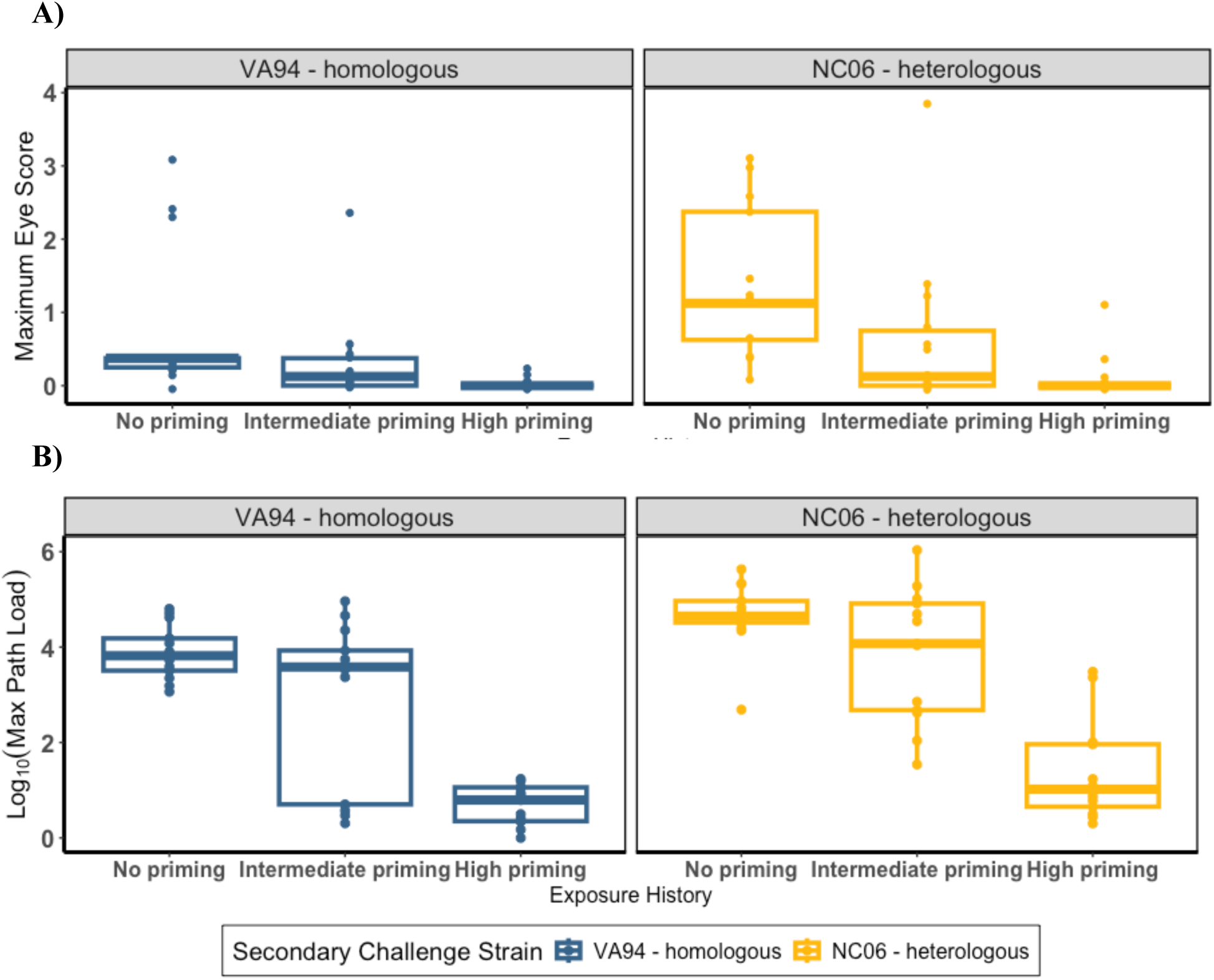
A) Maximum eye scores and B) pathogen loads after secondary challenge with one of two *Mycoplasma gallisepticum* strains (Left: homologous VA94, blue; Right: heterologous NC06, yellow) in index birds with distinct pathogen priming histories (x-axis: none, intermediate, or high). Each point represents maximum responses for each individual over four sample points. While eye scores are visualized as continuous here, they were analyzed as ordinal factors to account for their non-continuous distribution.

Pathogen loads just prior to reinfection challenge (day −4) were slightly elevated in index birds that received high priming treatment (Fig S1), suggesting complete clearance of high-dose priming had not universally occurred by the time of reinfection challenge. Nonetheless, birds that received high priming reached lower maximum pathogen loads during reinfection challenge than did birds with intermediate or no priming, indicating that residual pathogen from priming treatments was outweighed by effects of the protection acquired from priming (GLM assuming gamma distribution [parameters on *inverse* scale]: intermediate priming: 0.049 ± 0.036; high priming: 0.616 ± 0.10; priming treatment LR = 68.42, df = 2, P <0.0001; Fig. 2B). Priming treatment also interacted with strain identity to predict pathogen loads during reinfection (priming:secondary strain LR = 11.68, df = 2, P = 0.0023). Specifically, in index birds given high priming, hosts challenged with the heterologous, more virulent strain NC06 harbored significantly higher maximum pathogen loads than those challenged with VA94 (Table 1). Secondary strain did not have a significant main effect on pathogen loads (secondary strain LR = 0.245, df = 2, P = 0.62).

**Table 1.** Pairwise post-hoc comparisons for the significant interactive effect of priming (“prim.”) treatment (none, intermediate=”int”, or high) and secondary strain (VA94 or NC06) on maximum pathogen loads following secondary challenge. Interactive comparisons (strain differences within priming treatment) are in italics for ease of visualization. Upper triangle (light gray cells) gives Tukey-adjusted p-values (*\*p<0.05, **p<0.01*), the bottom triangle (unfilled cells) shows estimates for each pairwise comparison, and diagonal cells (dark gray, white text) show predicted pathogen loads (emmeans) for each combination on the response scale. Within the high priming treatment only, birds reinfected with NC06 had significantly higher pathogen loads than birds infected with VA94.

| *Priming: strain* | No priming<br>VA94 | Int. Prim.<br>VA94 | High Prim.<br>VA94 | No priming<br>NC06 | Int. prim.<br>NC06 | High prim.<br>NC06 |
| --- | --- | --- | --- | --- | --- | --- |
| No priming VA94 | 5.37 | 0.735 | <0.001** | 0.996 | 1.000 | 0.0007** |
| Intermediate VA94 | -0.049 | 4.24 | <0.001** | 0.436 | 0.656 | 0.0218* |
| High priming VA94 | -0.616 | -0.567 | 1.25 | <0.001** | <0.0001** | 0.0119* |
| No priming NC06 | 0.015 | 0.064 | 0.631 | 5.83 | 0.999 | 0.0002** |
| Intermediate NC06 | 0.004 | 0.054 | 0.620 | -0.011 | 5.49 | 0.0005** |
| High priming NC06 | -0.234 | -0.185 | 0.382 | -0.249 | -0.239 | 2.38 |

### Transmission Success

Pairwise transmission success was significantly lower, relative to unprimed birds, in reinfected index birds that received intermediate or high priming prior to secondary challenge, with the strongest reduction in transmission potential in the high priming group (logistic regression with a binomial distribution: intermediate priming: −0.7703 ± 0.381; high priming: − 1.581 ± 0.470; LR = 13.63, df = 2, P = 0.001; Fig 3). The heterologous, higher virulence NC06 strain had higher overall transmission success than the VA94 strain (NC06 strain: 0.8627 ± 0.351; LR = 6.45. df = 1, P = 0.011; Fig. 3). Although NC06 was notably the only strain with detectable transmission from high priming index birds, the low overall transmission success from hosts with high priming (2/13 pairs for NC06, relative to 0/13 for VA94) made any statistical interactions between secondary strain identity and priming treatment challenging to estimate.

**Figure 3.**
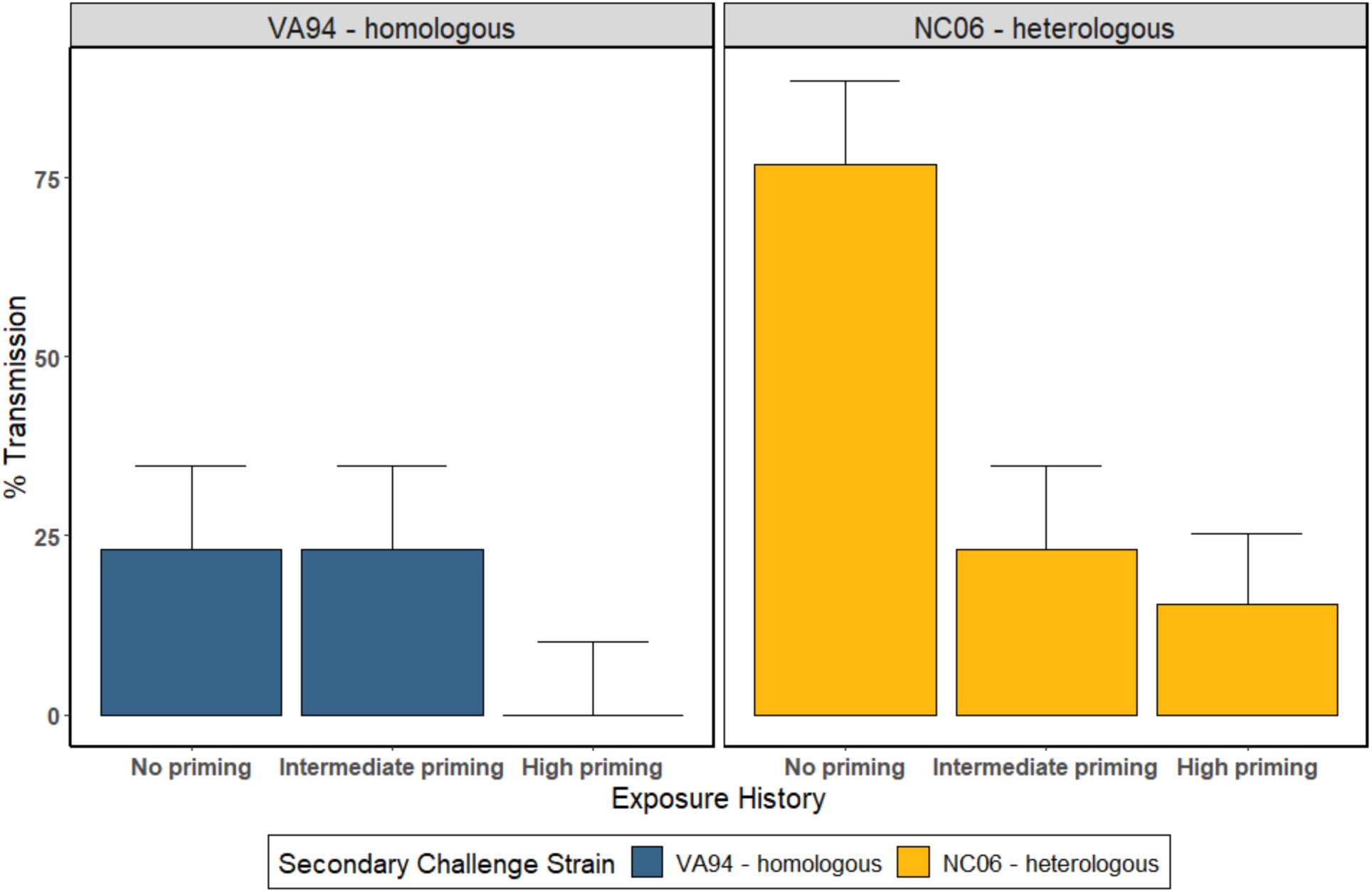
Pairwise transmission of *Mycoplasma gallisepticum* from captive house finches with variable pathogen priming (x-axis) that were experimentally inoculated with one of two pathogen strains (Left: VA94 - homologous, seen in blue; Right: NC06 - heterologous, seen in yellow). Percent transmission (y-axis) from index birds (transmitters) was measured in pathogen-naive cagemates. Higher degrees of priming reduced an index bird’s transmission potential to its cagemate for both strains, though the NC06 strain produced higher transmission regardless of host priming. Each treatment group had 13 pairs. Error bars represent binomial standard errors calculated using sample size.

### Host Factors Predictive of Transmission

The model of index bird transmission (Y/N) with the strongest support included an index bird’s maximum pathogen loads during secondary challenge, as well as the interaction between index bird maximum eye score and priming treatment. Specifically, across all priming treatments, index bird pathogen loads were positively associated with transmission probability (logistic regression: pathogen load: 3.93 ± 1.52, LR = 14.58, df = 1, P <0.001). In contrast, the relationship between an index bird’s maximum eye score and transmission success depended on a bird’s priming treatment (eye score*priming treatment: LR = 6.70, df = 2, P = 0.035; interaction visualized in Fig. 4). For birds given intermediate priming, higher maximum eye scores during reinfection were associated with slightly lower transmission (intermediate priming*score: −0.938 ± 0.601), while a positive effect of eye score was estimated for high primed birds (high priming*score: 46.1 ± 5055.7), although the model parameter could not be estimated with any precision due to the low number of high-primed birds with detectable eye score. The three-way interaction between pathogen load, disease severity, and priming treatment was not significant (LR = 0.44, df = 2, P = 0.80) and was removed from the model.

**Figure 4.**
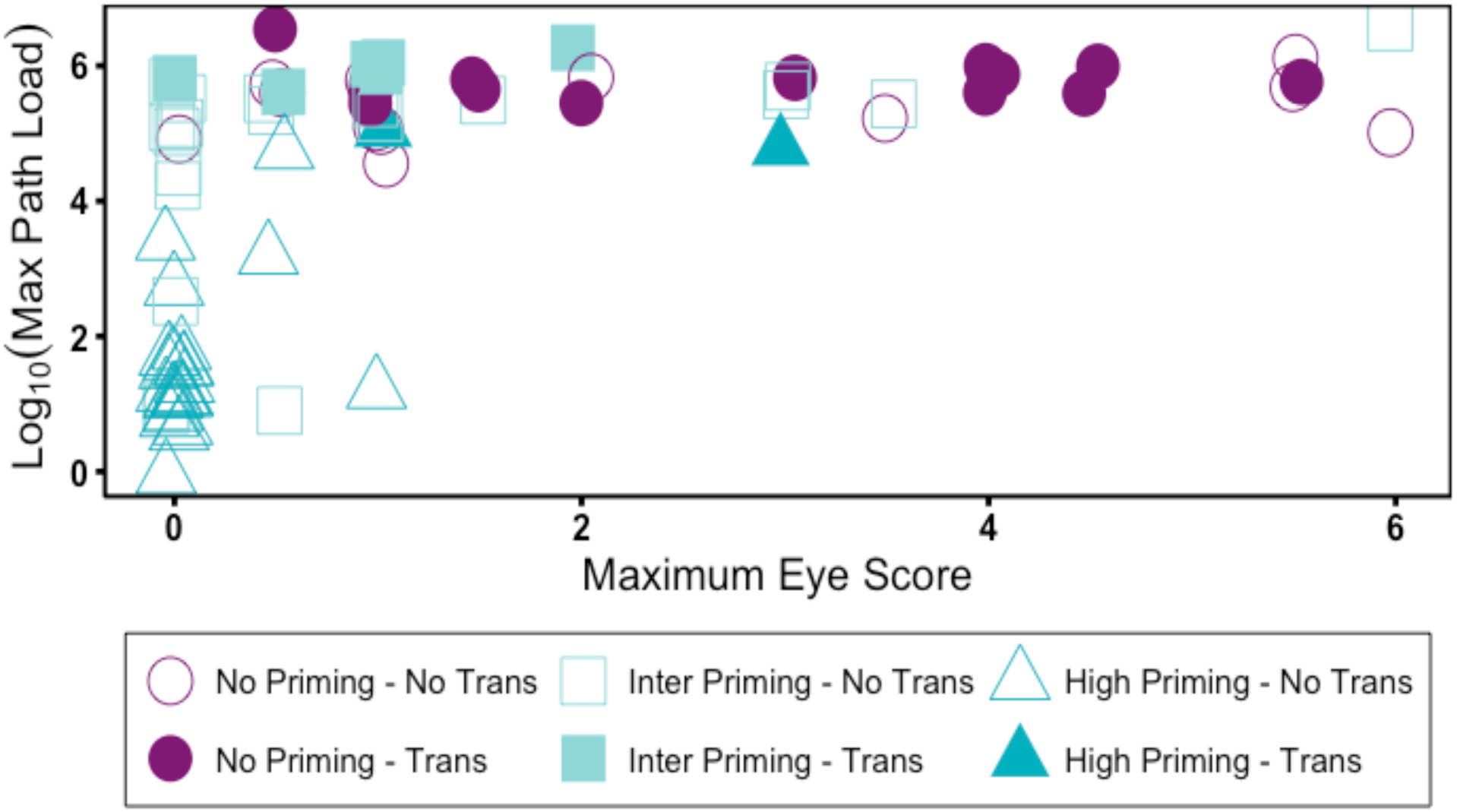
Successful pairwise transmission of *Mycoplasma gallisepticum* (filled symbols) from index house finches (n=78) given various priming treatments (symbols: unprimed birds = circles, intermediate priming = squares, high priming = triangles) was a function of the pathogen load of the index bird (y-axis), with successful transmission occurring only from index birds above a threshold pathogen load of log_10_ ∼ 4.5. There were also significant effects of index bird maximum eye score (x-axis) in interaction with pathogen priming treatment, with higher eye scores associated with lower transmission success at intermediate priming levels (squares), while for high priming birds (triangle), intermediate eye scores, which were the highest maximum reached for this treatment, were positively associated with transmission likelihood.

## Discussion

We show that pathogen priming significantly reduces transmission success during reinfections relative to birds that are infected for the first time. Nonetheless, reinfected hosts were still associated with a notable degree of successful pairwise transmission, particularly in cases where index birds had an intermediate priming level and reinfections were with a heterologous, higher virulence MG strain. Together, our findings suggest that reinfections meaningfully contribute to transmission dynamics in this system, particularly in cases that approximate our intermediate priming treatment, such as when pathogen priming in a population is variable, or acquired protection from previous infection has waned over time (37). As such, variation in the degree of acquired protection among hosts in natural populations, in systems where reinfections are common, can be an important impediment to transmission, placing selective pressure on invading pathogen strains to overcome host acquired protection.

Our main objective was to determine how the degree of pathogen priming, and thus the degree of acquired protection, alters host transmission potential during reinfection. For both strains examined, we found that the highest priming level resulted in strong reductions in transmission potential relative to unprimed hosts. These results are consistent with effects of host immunization on transmission likelihood in other key systems where this question has been examined, including mouse models (14, 15), and malaria (28) and SARS-Cov-2 in humans (27, 56–58). Together, these results demonstrate that prior pathogen infection or vaccination at levels sufficient to generate acquired protection can have strong effects on epidemic dynamics in wild populations, as documented in vaccinated human populations (e.g. 59), and likely provides strong selection on pathogens circulating in host populations where many individuals have acquired protection (17).

Our results also suggest that the degree of pathogen priming a host experienced has key effects on transmission success, potentially akin to the additive effects of vaccination and prior natural infection on the infectiousness of breakthrough SARS-Cov2 infections (27). Here, while our highest priming treatment caused strong reductions in transmission success for both strains, there were apparent, though not statistically significant, differences between the two strains in response to our intermediate priming level. The homologous VA94 strain showed no difference in transmission success between the no priming and intermediate priming levels. In contrast, the intermediate priming treatment resulted in fewer successful transmission events relative to unprimed birds for the heterologous NC06 strain. Future work should incorporate a low priming treatment such as a single, low-dose priming dose (39), to test whether there are strain-specific effects of lower degrees of pathogen priming on transmission in this system.

We also detected overall strain differences in transmission success akin to prior work (41), with the NC06 strain showing higher transmission success than VA94, regardless of a host’s priming treatment. For within-host pathogen loads, however, effects of strain were dependent on host priming background, with the NC06 strain reaching significantly higher within-host pathogen loads than the VA94 strain within the high-priming group. This pathogen load advantage for NC06 may explain why there was successful transmission (∼15%) of the NC06 strain even from hosts with the high priming level, whereas there were no instances of successful transmission of the VA94 strain from hosts with the same high priming level. Overall, our findings align with mouse-malaria studies showing that parasite clone differences in growth rate and transmission are largely maintained in immunized hosts, despite overall reductions in average growth and transmission (15). Although heterology of the reinfection strain relative to the priming strain may partly explain the ability of NC06 to successfully replicate within and transmit during reinfection from hosts with the highest priming level, heterology is unlikely to be the main driver of these patterns: first, heterology is only a relevant mechanism in primed hosts, whereas transmission success of NC06 was substantially higher in unprimed hosts, for which the NC06 strain was the first MG strain that such hosts had been exposed to (and thus was not truly “heterologous”). Second, if heterology was an important driver of transmission potential during reinfection, we would predict that NC06 strain transmission would have been less affected by our intermediate priming level than was the homologous strain; instead NC06 appears more strongly affected (relative to unprimed birds) by intermediate priming than was VA94, the homologous strain. Finally, prior work in this system using larger numbers of MG strains found that strain virulence was a stronger predictor of reinfection likelihood than homology (8, 40), although effects of homology may result in small increases in acquired protection (8). Overall, our results complement prior work showing higher transmission rates (41, 44) for virulent MG strains in immunologically-naive hosts, and suggest that this transmission advantage is maintained in the presence of strong acquired protection from priming.

We can use our results to hypothesize as to how detected effects of pathogen priming on transmission potential might ultimately influence virulence evolution in this system, although studies using replicate heterologous strains are needed to confirm effects of strain virulence on transmission potential during reinfection. For directly-transmitted pathogens, where virulence typically reduces opportunities for transmission (60), high virulence should be favored when it is an unavoidable by-product of the high within-host replication needed for transmission given a contact (20). Thus, any within-host selective environment that requires higher levels of replication to achieve transmission should favor higher pathogen virulence, a prediction supported by prior studies in the mouse-malaria system (22, 23). Because pathogen priming led to significant reductions in pathogen loads and transmission success in reinfected hosts in our study, and maximum pathogen loads were associated with successful transmission, a host population with acquired protection from priming would favor a higher optimal pathogen virulence relative to an unprimed, naïve host population. Second, optimal virulence is predicted to increase whenever acquired protection in hosts reduces the costs of virulence to pathogens (8, 15, 17) by preventing reinfected hosts from dying when infected with strains of higher virulence. These predictions were confirmed empirically in chickens vaccinated with Marek’s virus (17), which are able to successfully transmit virulent strains in vaccinated flocks. Notably, priming at both intermediate and high levels in our study resulted in significantly lower index bird disease severity during reinfection, and disease severity is a relevant proxy for infection-mediated mortality in this system (61). Thus, while survival did not vary here (because birds survive MG infection in captive conditions where predators are absent), our results suggest that primed hosts in the wild are more likely to survive reinfections, potentially resulting in longer infectious periods for virulent strains in primed hosts (17). Overall, our findings, alongside prior modeling work on this system (8), suggest that acquired protection from priming in this system will favor higher optimal pathogen virulence.

Our results also reveal a key role for pathogen load in driving transmission success among individuals. In contrast to work by Bonneaud at al. (44) that did not detect effects of strain-level variation in pathogen loads on MG transmission success, we found that individual variation in pathogen loads was positively associated with pairwise transmission success, regardless of host priming background. Although we also predicted that individual variation in disease severity would correlate with transmission as seen previously in this system (44, 45), we found that associations of disease severity with transmission success depended on a bird’s priming treatment. For intermediate-primed birds, higher disease severity was associated with *reductions* rather than increases in transmission success, such that several birds with both high levels of disease severity and pathogen load did not successfully transmit to cagemates. While the reasons for this are unclear, one possibility is that because birds with high levels of disease severity also show behavioral morbidity during infection (61), high levels of disease severity may sometimes suppress rather than augment transmission opportunities in this system, and likely others (62). Indeed, prior work (45) found that MG-infected house finches that maintained foraging activity were more likely to transmit to cagemates, consistent with the idea that behavioral morbidity can suppress transmission. In contrast, for high-primed birds, disease severity appeared positively associated with transmission success, but due to the strong effects of high priming on protection from disease, we did not have sufficient numbers of high-primed birds with detectable eye scores to estimate this relationship with any precision. Overall, our findings suggest that effects of disease severity on transmission success differ between reinfected and naive hosts, with potential consequences for virulence evolution.

Overall, our results demonstrate that acquired protection from pathogen priming alters transmission success, potentially in distinct ways for different pathogen strains. Our results add to broader growing recognition (9, 12), that many epidemiological systems fall between the classically studied Susceptible-Infected-Recovered (SIR) systems, where recovered individuals acquire complete protection that can wane over time, and Susceptible-Infected-Susceptible (SIS) systems, where hosts acquire no lasting protection from infection. As mathematical models continue to evolve to better capture this variation in infection- or vaccine-induced immunity (9, 63, 64), it is important to empirically quantify how parameters such as transmission success vary with acquired protection in both vaccinated and reinfected populations to better understand transmission dynamics and predict the evolution of more harmful pathogens.

## Acknowledgments

This work was funded by NIH grants R01GM105245 and R01GM144972 as part of the joint NIH-NSF-USDA Ecology and Evolution of Infectious Diseases program. Additional fellowship support for A. Leon was provided by the Virginia Tech IMSD program funded through NIH-NIGMS grant R25GM072767-09, the Virginia Tech College of Science Graduate Departmental Fellowship Award through the Department of Biological Sciences, the College of Science Roundtable Scholarship and the Southern Regional Education Board’s Dissertation Writing Fellowship. We thank Courtney Thomason, Sahnzi Moyers, and Matt Aberle for their assistance with this project, as well as Kate Langwig, Joel McGlothlin, Rami Dalloul and Liwu Li for valuable feedback. We especially thank Natalie Bale and Eddie Schuler for their help with data collection, as well as Daphne Aguirre, Jennifer Holub and Camron Robertson.

## Supplemental Materials

**Figure S1.**
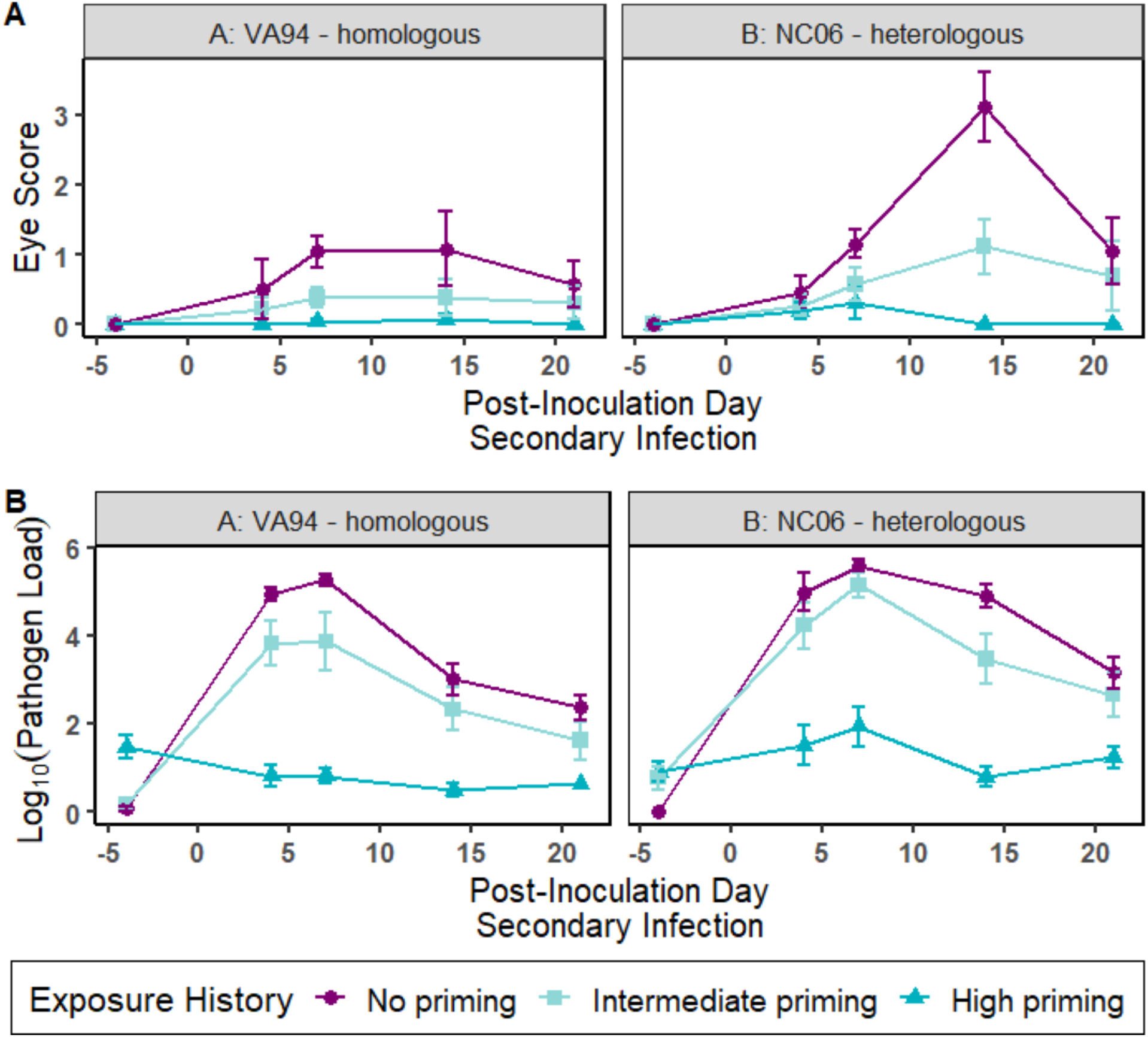
A) Disease severity (eye score) and B) pathogen loads over the course of secondary infection for index birds with distinct levels of pathogen priming given a secondary high-dose challenge with one of two strains of *Mycoplasma gallisepticum* (A: VA94; B: NC06). Exposure history (purple, circles – no priming; light blue, squares – intermediate priming; dark blue, triangular points – high priming) largely determined the extent of disease and pathogen load across both strains, although the heterologous strain (NC06) produced higher levels of disease than the homologous strain (VA94), overall. Error bars represent standard error from the mean.

### Bird Capture and Housing

Hatch-year house finches were captured in Montgomery County, VA in June-July 2016 using a combination of mesh wire traps and mist-nets under permits from VDGIF (056090) and USFWS (MB158404-1). To ensure that birds used in our experiments had no previous exposure to MG in the wild, captive animals underwent a two-week quarantine protocol wherein they were monitored for visible signs of infection and blood sampled on day 14 post-capture to test for MG-specific antibodies (as per (65)). Only individuals that never showed clinical signs of infection, had not been housed with an infected individual, and were seronegative for pathogen-specific antibodies were included in the experiment (n=156 total).

All animals were pair-housed during quarantine, but index birds were single-housed prior to the start of and for the duration of the priming portion of the experiment. After recovery from priming exposures and immediately following secondary challenge, the index birds were then pair-housed with MG-naïve cagemates to assess pairwise transmission potential during reinfection. For the entirety of their time in captivity, finches were held at constant day length (12L:12D) and temperature, and were fed an *ad libitum* diet (Daily Maintenance Diet, Roudybush Inc., Woodland, CA). Individuals were given food in open-cup dishes for the priming portion of the study (when no transmission could occur due to individual housing). When inoculated birds were pair-housed with pathogen-naïve cagemates to quantify pairwise transmission, all pairs were given a two-port hanging tube feeder to mimic the feeder type most likely to facilitate transmission in the wild (34, 66).

### Inoculation

Stock inocula were grown in Frey’s broth media with 15% swine serum (FMS) and provided by D.H. Ley, North Carolina State University, College of Veterinary Medicine, Raleigh, NC, USA. All inocula were stored at −80°C and thawed and diluted immediately before use. Inoculation dilutions for priming exposures were calculated using the starting viable count of 10^7^ CCU/ml of VA94. To control for the stress of extra handling and inoculation for birds in the repeated low-dose priming group, a randomly selected subset of individuals from the no priming and high-dose priming groups were given a sham inoculation of sterile FMS on priming days 1, 3, 5, 7 and 9 (Fig. 1). No effect of this sham treatment was detected on disease (F = 0.048, df = 30, P = 0.83) or infection outcomes (F = 1.28, df = 12, P = 0.28) compared to control animals not given sham inoculations.

### Pathogen load quantification

Both conjunctival sacs were swabbed for 5 seconds using separate sterile cotton swabs dipped in tryptose phosphate broth (TPB) and eluted in a single tube containing 300uL of TPB. Samples were kept on ice until frozen at −20°C and remained frozen until thawed for DNA extraction. DNA was extracted using Qiagen DNeasy 96 Blood and Tissue kits (Qiagen, Valencia, CA). Quantitative polymerase chain reaction (qPCR) was performed using a Bio-Rad C1000 CFX96 Real-time System (Hercules, CA). Primers and probes that target the Mgc2 gene of MG were used, and a standard curve of 2.98 x 10^1^ to 2.98 x 10^8^ copy numbers was produced using a plasmid containing a 303 bp Mgc2 insert (67). Cycling parameters used were as follows: 95°C for 3 minutes then 40 cycles of 95°C for 3 seconds followed by 60°C for 30 seconds.

## References Cited

1. Yamamoto T, Nagasawa I, Nojima M, Yoshida K, Kuwabara Y. 1999. Sexual transmission and reinfection of group B streptococci between spouses. J Obstet Gynaecol Res 25:215–219.

2. Nardell E, McInnis B, Thomas B, Weidhaas S. 1986. Exogenous reinfection with tuberculosis in a shelter for the homeless. N Engl J Med 315:1570–1575.

3. Islam N, Krajden M, Shoveller J, Gustafson P, Gilbert M, Buxton JA, Wong J, Tyndall MW, Janjua NZ, British Columbia Hepatitis Testers Cohort (BC-HTC) team. 2017. Incidence, risk factors, and prevention of hepatitis C reinfection: a population-based cohort study. Lancet Gastroenterol Hepatol 2:200–210.

4. Versteegh FGA, Schellekens JFP, Nagelkerke AF, Roord JJ. 2007. Laboratory-confirmed reinfections with Bordetella pertussis. Acta Paediatr 91:95–97.

5. Pilz S, Theiler-Schwetz V, Trummer C, Krause R, Ioannidis JPA. 2022. SARS-CoV-2 reinfections: Overview of efficacy and duration of natural and hybrid immunity. Environ Res 209:112911.

6. Netea MG, Schlitzer A, Placek K, Joosten LAB, Schultze JL. 2019. Innate and Adaptive Immune Memory: an Evolutionary Continuum in the Host’s Response to Pathogens. Cell Host Microbe 25:13–26.

7. Forshey BM, Reiner RC, Olkowski S, Morrison AC, Espinoza A, Long KC, Vilcarromero S, Casanova W, Wearing HJ, Halsey ES, Kochel TJ, Scott TW, Stoddard ST. 2016. Incomplete Protection against Dengue Virus Type 2 Re-infection in Peru. PLoS Negl Trop Dis 10:e0004398.

8. Fleming-Davies AE, Williams PD, Dhondt AA, Dobson AP, Hochachka WM, Leon AE, Ley DH, Osnas EE, Hawley DM. 2018. Incomplete host immunity favors the evolution of virulence in an emergent pathogen. Science 359:1030–1033.

9. Le A, King AA, Magpantay FMG, Mesbahi A, Rohani P. 2021. The impact of infection-derived immunity on disease dynamics. J Math Biol 83:61.

10. Breathnach AS, Riley PA, Cotter MP, Houston AC, Habibi MS, Planche TD. 2021. Prior COVID-19 significantly reduces the risk of subsequent infection, but reinfections are seen after eight months. J Infect 82:e11–e12.

11. Goldberg Y, Mandel M, Bar-On YM, Bodenheimer O, Freedman LS, Ash N, Alroy-Preis S, Huppert A, Milo R. 2022. Protection and Waning of Natural and Hybrid Immunity to SARS-CoV-2. N Engl J Med 386:2201–2212.

12. Gomes MGM, White LJ, Medley GF. 2004. Infection, reinfection, and vaccination under suboptimal immune protection: epidemiological perspectives. J Theor Biol 228:539–549.

13. Raida MK, Buchmann K. 2009. Innate immune response in rainbow trout (Oncorhynchus mykiss) against primary and secondary infections with Yersinia ruckeri O1. Dev Comp Immunol 33:35–45.

14. Buckling A, Read AF. 2001. The effect of partial host immunity on the transmission of malaria parasites. Proc Biol Sci 268:2325–2330.

15. Mackinnon MJ, Read AF. 2003. The effects of host immunity on virulence–transmissibility relationships in the rodent malaria parasite Plasmodium chabaudi. Parasitology 126:103–112.

16. Singanayagam, Hakki, Dunning. Community transmission and viral load kinetics of the SARS-CoV-2 delta (B. 1.617. 2) variant in vaccinated and unvaccinated individuals in the UK: a …. Lancet Infect Dis.

17. Read AF, Baigent SJ, Powers C, Kgosana LB, Blackwell L, Smith LP, Kennedy DA, Walkden-Brown SW, Nair VK. 2015. Imperfect Vaccination Can Enhance the Transmission of Highly Virulent Pathogens. PLoS Biol 13:e1002198.

18. Chase-Topping ME, Pooley C, Moghadam HK, Hillestad B, Lillehammer M, Sveen L, Doeschl-Wilson A. 2021. Impact of vaccination and selective breeding on the transmission of Infectious salmon anemia virus. Aquaculture 535:736365.

19. Chase-Topping M, Xie J, Pooley C, Trus I, Bonckaert C, Rediger K, Bailey RI, Brown H, Bitsouni V, Barrio MB, Gueguen S, Nauwynck H, Doeschl-Wilson A. 2020. New insights about vaccine effectiveness: Impact of attenuated PRRS-strain vaccination on heterologous strain transmission. Vaccine 38:3050–3061.

20. Acevedo MA, Dillemuth FP, Flick AJ, Faldyn MJ, Elderd BD. 2019. Virulence-driven trade-offs in disease transmission: A meta-analysis. Evolution 73:636–647.

21. Lipsitch M, Moxon ER. 1997. Virulence and transmissibility of pathogens: what is the relationship? Trends Microbiol 5:31–37.

22. Barclay VC, Sim D, Chan BHK, Nell LA, Rabaa MA, Bell AS, Anders RF, Read AF. 2012. The evolutionary consequences of blood-stage vaccination on the rodent malaria Plasmodium chabaudi. PLoS Biol 10:e1001368.

23. Mackinnon MJ, Read AF. 2004. Immunity promotes virulence evolution in a malaria model. PLoS Biol 2:E230.

24. Bekliz M, Adea K, Vetter P, Eberhardt CS, Hosszu-Fellous K, Vu D-L, Puhach O, Essaidi-Laziosi M, Waldvogel-Abramowski S, Stephan C, L’Huillier AG, Siegrist C-A, Didierlaurent AM, Kaiser L, Meyer B, Eckerle I. 2022. Neutralization capacity of antibodies elicited through homologous or heterologous infection or vaccination against SARS-CoV-2 VOCs. Nat Commun 13:3840.

25. Markov PV, Katzourakis A, Stilianakis NI. 2022. Antigenic evolution will lead to new SARS-CoV-2 variants with unpredictable severity. Nat Rev Microbiol 20:251–252.

26. Mideo N, Kamiya T. 2022. Antigenic evolution can drive virulence evolution. Nat Ecol Evol 6:24– 25.

27. Tan ST, Kwan AT, Rodríguez-Barraquer I, Singer BJ, Park HJ, Lewnard JA, Sears D, Lo NC. 2023. Infectiousness of SARS-CoV-2 breakthrough infections and reinfections during the Omicron wave. Nat Med 29:358–365.

28. Ranawaka MB, Munesinghe YD, de Silva DM, Carter R, Mendis KN. 1988. Boosting of transmission-blocking immunity during natural Plasmodium vivax infections in humans depends upon frequent reinfection. Infect Immun 56:1820–1824.

29. Konrad M, Vyleta ML, Theis FJ, Stock M, Tragust S, Klatt M, Drescher V, Marr C, Ugelvig LV, Cremer S. 2012. Social transfer of pathogenic fungus promotes active immunisation in ant colonies. PLoS Biol 10:e1001300.

30. Müller-Klein N, Risely A, Schmid DW, Manser M, Clutton-Brock T, Sommer S. 2022. Two decades of tuberculosis surveillance reveal disease spread, high levels of exposure and mortality and marked variation in disease progression in wild meerkats. Transbound Emerg Dis 69:3274–3284.

31. Weitzman CL, Ceja G, Leon AE, Hawley DM. 2022. Protection Generated by Prior Exposure to Pathogens Depends on both Priming and Challenge Dose. Infection and Immunity 10.1128/iai.00537-21.

32. Glover M, Colombo SAP, Thornton DJ, Grencis RK. 2019. Trickle infection and immunity to Trichuris muris. PLoS Pathog 15:e1007926.

33. Dhondt AA, Altizer S, Cooch EG, Davis AK, Dobson A, Driscoll MJL, Hartup BK, Hawley DM, Hochachka WM, Hosseini PR, Jennelle CS, Kollias GV, Ley DH, Swarthout ECH, Sydenstricker KV. 2005. Dynamics of a novel pathogen in an avian host: Mycoplasmal conjunctivitis in house finches. Acta Trop 94:77–93.

34. Adelman JS, Moyers SC, Farine DR, Hawley DM. 2015. Feeder use predicts both acquisition and transmission of a contagious pathogen in a North American songbird. Proceedings of the Royal Society B: Biological Sciences 282:20151429.

35. Dhondt AA, Dhondt KV, Hawley DM, Jennelle CS. 2007. Experimental evidence for transmission of Mycoplasma gallisepticum in house finches by fomites. Avian Pathol 36:205–208.

36. Faustino CR, Jennelle CS, Connolly V, Davis AK, Swarthout EC, Dhondt AA, Cooch EG. 2004. Mycoplasma gallisepticum infection dynamics in a house finch population: seasonal variation in survival, encounter and transmission rate. J Anim Ecol 73:651–669.

37. Sydenstricker KV, Dhondt AA, Ley DH, Kollias GV. 2005. Re-exposure of captive house finches that recovered from Mycoplasma gallisepticum infection. J Wildl Dis 41:326–333.

38. Dhondt AA, Dhondt KV, Hochachka WM, Ley DH, Hawley DM. 2017. Response of House Finches Recovered from Mycoplasma gallisepticum to Reinfection with a Heterologous Strain. Avian Dis 61:437–441.

39. Leon AE, Hawley DM. 2017. Host Responses to Pathogen Priming in a Natural Songbird Host. Ecohealth 14:793–804.

40. Leon AE, Fleming-Davies AE, Hawley DM. 2019. Host exposure history modulates the within-host advantage of virulence in a songbird-bacterium system. Sci Rep 9:20348.

41. Williams PD, Dobson AP, Dhondt KV, Hawley DM, Dhondt AA. 2014. Evidence of trade-offs shaping virulence evolution in an emerging wildlife pathogen. J Evol Biol 27:1271–1278.

42. Adelman JS, Carter AW, Hopkins WA, Hawley DM. 2013. Deposition of pathogenic Mycoplasma gallisepticum onto bird feeders: host pathology is more important than temperature-driven increases in food intake. Biol Lett 9:20130594.

43. Hawley DM, Thomason CA, Aberle MA, Brown R, Adelman JS. 2023. High virulence is associated with pathogen spreadability in a songbird–bacterial system. Royal Society Open Science 10:220975.

44. Bonneaud C, Tardy L, Hill GE, McGraw KJ, Wilson AJ, Giraudeau M. 2020. Experimental evidence for stabilizing selection on virulence in a bacterial pathogen. Evol Lett 4:491–501.

45. Ruden RM, Adelman JS. 2021. Disease tolerance alters host competence in a wild songbird. Biol Lett 17:20210362.

46. Hawley, DM, Thomason C, Aberle M, Brown R, and Adelman JS. High virulence is associated with pathogen spreadability in a songbird-bacterial system.

47. Hawley DM, Osnas EE, Dobson AP, Hochachka WM, Ley DH, Dhondt AA. 2013. Parallel patterns of increased virulence in a recently emerged wildlife pathogen. PLoS Biol 11:e1001570.

48. Bonneaud C, Giraudeau M, Tardy L, Staley M, Hill GE, McGraw KJ. 2018. Rapid Antagonistic Coevolution in an Emerging Pathogen and Its Vertebrate Host. Curr Biol 28:2978–2983.e5.

49. Ley DH, Edward Berkhoff J, McLaren JM. 1996. Mycoplasma gallisepticum Isolated from House Finches (Carpodacus mexicanus) with Conjunctivitis. Avian Diseases 10.2307/1592250.

50. Sydenstricker KV, Dhondt AA, Hawley DM, Jennelle CS, Kollias HW, Kollias GV. 2006. Characterization of experimental Mycoplasma gallisepticum infection in captive house finch flocks. Avian Dis 50:39–44.

51. R Development Core Team. 2020. R: A Language and Environment for Statistical Computing (4.0.3).

52. Fox, Weisberg, Adler, Bates. Package “car.” : R Foundation for ….

53. Christensen RHB. 2015. ordinal—regression models for ordinal data. R package version 28:2015.

54. Bates D, Mächler M, Bolker B, Walker S. 2014. Fitting Linear Mixed-Effects Models using lme4. arXiv [statCO].

55. Dormann CF, Elith J, Bacher S, Buchmann C, Carl G, Carré G, Marquéz JRG, Gruber B, Lafourcade B, Leitão PJ, Münkemüller T, McClean C, Osborne PE, Reineking B, Schröder B, Skidmore AK, Zurell D, Lautenbach S. 2013. Collinearity: a review of methods to deal with it and a simulation study evaluating their performance. Ecography 36:27–46.

56. Abrokwa SK, Müller SA, Méndez-Brito A, Hanefeld J, El Bcheraoui C. 2021. Recurrent SARS-CoV-2 infections and their potential risk to public health - a systematic review. PLoS One 16:e0261221.

57. Puhach O, Adea K, Hulo N, Sattonnet P, Genecand C, Iten A, Jacquérioz F, Kaiser L, Vetter P, Eckerle I, Meyer B. 2022. Infectious viral load in unvaccinated and vaccinated individuals infected with ancestral, Delta or Omicron SARS-CoV-2. Nat Med 28:1491–1500.

58. Bailey RI, Cheng HH, Chase-Topping M, Mays JK, Anacleto O, Dunn JR, Doeschl-Wilson A. Pathogen transmission from vaccinated hosts can cause dose-dependent reduction in virulence 10.1101/830570.

59. Anderson RM, May RM. 1985. Vaccination and herd immunity to infectious diseases. Nature 318:323–329.

60. Anderson RM, May RM. 1982. Coevolution of hosts and parasites. Parasitology 85 (Pt 2):411–426.

61. Adelman JS, Mayer C, Hawley DM. 2017. Infection reduces anti-predator behaviors in house finches. J Avian Biol 48:519–528.

62. Kennedy DA. 2023. Death is overrated: the potential role of detection in driving virulence evolution. Proc Biol Sci 290:20230117.

63. Katriel G. 2010. Epidemics with partial immunity to reinfection. Math Biosci 228:153–159.

64. Magpantay FMG, Riolo MA, DE Cellès MD, King AA, Rohani P. 2014. EPIDEMIOLOGICAL CONSEQUENCES OF IMPERFECT VACCINES FOR IMMUNIZING INFECTIONS. SIAM J Appl Math 74:1810–1830.

65. Hawley DM, Grodio J, Frasca S, Kirkpatrick L, Ley DH. 2011. Experimental infection of domestic canaries (Serinus canaria domestica) with Mycoplasma gallisepticum: a new model system for a wildlife disease. Avian Pathol 40:321–327.

66. Hartup BK, Mohammed HO, Kollias GV, Dhondt AA. 1998. Risk factors associated with mycoplasmal conjunctivitis in house finches. J Wildl Dis 34:281–288.

67. Grodio JL, Dhondt KV, O’Connell PH, Schat KA. 2008. Detection and quantification of Mycoplasma gallisepticum genome load in conjunctival samples of experimentally infected house finches (Carpodacus mexicanus) using real-time polymerase chain reaction. Avian Pathol 37:385– 391.

